# COVIDOUTCOME – Estimating COVID Severity Based on Mutation Signatures in the SARS-CoV-2 Genome

**DOI:** 10.1101/2021.04.01.438063

**Authors:** Ádám Nagy, Balázs Ligeti, János Szebeni, Sándor Pongor, Balázs Győrffy

## Abstract

**Introduction:** Numerous studies demonstrate frequent mutations in the genome of SARS-CoV-2. Our goal was to statistically link mutations to severe disease outcome.

**Methods:** We used an automated machine learning approach where 1,594 viral genomes with available clinical follow-up data were used as the training set (797 “severe” and 797 “mild”). The best algorithm, based on random forest classification combined with the LASSO feature selection algorithm was employed to the training set to link mutation signatures and outcome. The performance of the final model was estimated by repeated, stratified, 10-fold cross validation (CV), then adjusted for multiple testing with Bootstrap Bias Corrected CV.

**Results:** We identified 26 protein and UTR mutations significantly linked to severe outcome. The best classification algorithm uses a mutation signature of 22 mutations as well as the patient’s age as the input and shows high classification efficiency with an AUC of 0.94 (CI: [0.912, 0.962]) and a prediction accuracy of 87% (CI: [0.830, 0.903]). Finally, we established an online platform (https://covidoutcome.com/) which is capable to use a viral sequence and the patient’s age as the input and provides a percentage estimation of disease severity.

**Discussion:** We demonstrate a statistical association between mutation signatures of SARS-CoV-2 and severe outcome of COVID-19. The established analysis platform enables a real-time analysis of new viral genomes.

**KEY MESSAGES:**

1. A statistical link between SARS-Cov-2 mutation status and severe COVID outcome was established using automated machine learning techniques based on random forest and logistic regression combined with feature selection algorithms.
2. A mutation signature based on 3,779 protein coding and 36 UTR mutations capable to identify severe outcome cases was established.
3. The trained model showed high classification performance (AUC=0.94 (CI: [0.912, 0.962]), accuracy=0.87 (CI: [0.830, 0.903])).
4. A registration-free web-server for automated classification of new samples was set up and is accessible at http://www.covidoutcome.com.
5. The established pipeline provides a quick assessment of future patients warranting a prospective clinical validation.

## INTRODUCTION

With several hundred thousand fully sequenced genomes deposited in various databases, coronavirus SARS-Cov-2, the causative agent of the COVID-19 pandemic is probably the most thoroughly sequenced organism today. The variance we see is impressive: there is no or hardly any sequence position in the genome that is not mutated in one of the published sequences.

Interpretation of SARS-Cov-2 genome data, especially in terms of disease severeness and patient mortality is a formidable task complicated by facts such as the virus spreading in a constantly mixing human population, in differentially susceptible age groups, and in vastly different healthcare conditions (e.g. (1)). In addition, only a small part of deposited genomes are annotated with patient status data. Consequently, one can argue that mutations are simply neutral regional markers that rarely affect viral fitness and clinical outcome. On the other hand, there is a growing body of empirical evidence showing that specific mutation patterns such as Spike protein mutation D614G and its accompanying mutations are associated with faster spreading of the virus (2), (3) and it was shown that Spike D614G mutants not only spread faster but also cause more severe disease in animal models (4). Recent statistical studies of about five thousand SARS-Cov-2 genome sequences showed that various mutations were significantly associated with clinical outcome and it was found that many of the mutations affected known functional parts of the Spike and Nucleocapsid proteins (5), (6). It is an open question whether or not the mutation signature of SARS-Cov-2 genomes can be used as an indicator of disease severity given the current data available.

Machine learning classification algorithms (such as support vector machines (7,8), random forest (9), logistic regression (10) among many others) are par excellence tools for uncovering hidden associations in large datasets. Given two sets of samples assigned to different classes (such as disease outcomes), and a mathematical description for the samples (such as a vector, or a list of mutations), classification algorithms can give well-understood statistical estimates regarding how well a mathematical description can discriminate the two classes. The pertinent measures are defined in the framework of ROC – receiver operating characteristics – analysis (11,12). On the other hand, mutation lists – that we term here mutation signatures – can be quite long and difficult to handle. Feature selection algorithms – such as the Lasso algorithm or the more recent Statistically Equivalent Signatures (SES) method (13), – can help one to condense a mutation list to an essential core set. And if such a recurrent set exists across various datasets and classification algorithms, one is encouraged to believe that there is an association between sample descriptions and the class definitions – in our case mutation signatures and disease outcomes.

Here we applied machine learning classification combined with feature selection algorithms to a cohort of 1594 SARS-Cov-2 genomes and their associated patient data in order to show that the known mutation signatures contain the information sufficient to separate mild and severe outcome classes and can be considered as predictors of severe outcomes. We also established an online analysis platform for predicting the probability of severe infection, starting from a SARS-Cov-2 genome sequence.

## MATERIALS AND METHODS

The SARS-CoV-2 nucleic acid sequences were downloaded in FASTA format from the GISAID virus repository (https://www.gisaid.org/, accessed on December 2, 2020). Only genome sequences annotated with patient follow-up status data were downloaded. The CoVsurver analysis tool (https://corona.bii.a-star.edu.sg) was used to extract the mutations. The viral sequences in FASTA format were used as input for this tool. The “hCoV-19/Wuhan/WIV04/2019” strain was used as the reference. The UTR mutations were extracted from the multiple alignment of underlying sequences by comparing the target sequence to the reference sequence. The multiple alignment was constructed using the MAFF software tool (14), and substitutions occurring in at least ten genomes were selected for further analysis. The protein mutations were exported in protein alteration format, non-protein (i.e. UTR) mutations were exported in nucleotide mutation format.

Artificial intelligence (machine learning) algorithms were used to identify mutations associated with the sever outcome. Briefly, we chose a procedure, using the JADBio platform (15), that starts with genomic mutation data as the input, carrying out classification based on rigde logistic regression (RLR) (10), random forests (9) or support vector machines (7) in conjunction with LASSO feature selection (16) whenever appropriate, and outputting a) classification efficiency measures (accuracy and AUC -for a review see (12)) and, b) a feature importance list i.e. a list of mutations ranked according to their importance in distinguishing severe vs. other outcomes. For training and testing the classification and feature selection algorithms, we organized the genome data into three datasets: Dataset #1 included 797 severe and 797 mild cases **(Supplemental Table 1)**; and Dataset #2 included 638 severe and 638 mild samples **(Supplemental Table 2)**.

The online analysis platform (https://www.covidoutcome.com) is written in R using the R Shiny package (https://CRAN.R-project.org/package=shiny) and is running under Linux Debian 64-bit (x86_64). The server takes a SARS-CoV-2 genomic sequence in FASTA format as the input, and provides a) a list of protein as well as UTR mutations and b) a probability of the input genome producing severe infection, as the output.

The server fist performs a global pairwise sequence alignment between the input sequence and the reference genome of the “Wuhan strain” (hCoV-19/Wuhan/WIV04/2019), using the program of the “Biostrings” R Bioconductor package (https://bioconductor.org/packages/Biostrings/), and outputs the protein as well as UTR mutations. The second step of the analysis includes prediction of the clinical outcome which is expressed as the probability of severe outcome. The prediction is based on the Random Forest model trained on 797 “mild” and 797 “severe” genome records (Dataset 1). The result is presented in numerical as well as graphical form. **Figure 1** shows the complete analysis workflow.

**Figure 1.**
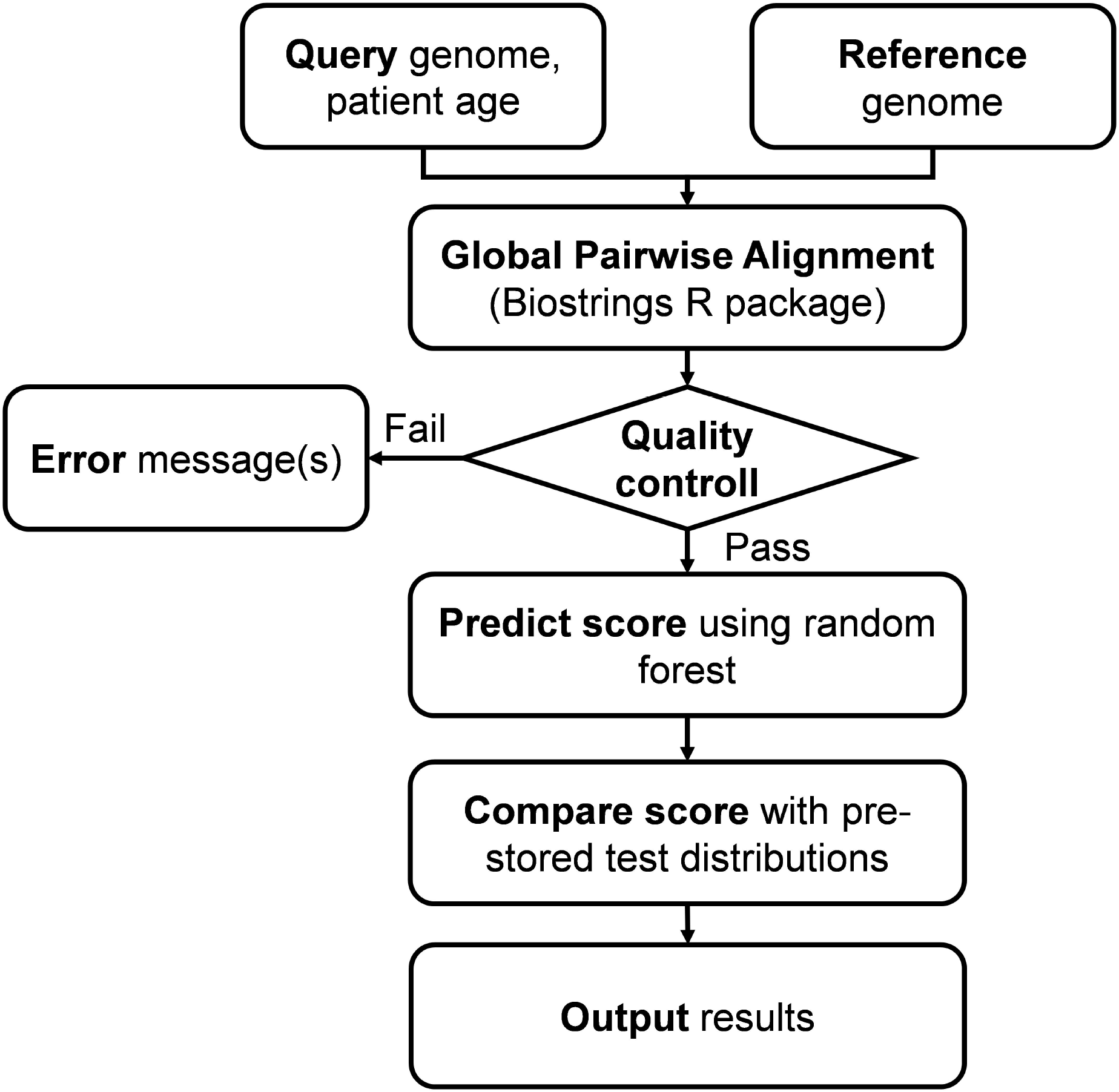
Flowchart of the online analysis platform. Quality control includes checking the number of identities with the Wuhan strain (min. 90%), genome length (29,000<length<40,000), GC contents (37%<GC<39%) and number of uncertain („N”) characters (max 2%).

## RESULTS

### Set up of viral datasets

We retrieved from the GISAID database a total of 9,781 SARS-CoV-2 genome data that were provided with patient status indications. We found that patient status was described with 179 different, submitter-defined terms, so we formed cohorts that included clearly defined patient descriptions. This was possible for the mildest and for the most severe outcomes that we designated as “mild” and “severe”. The pertinent terms are listed in **Supplemental Table 3**. Hospitalized patients were more difficult to categorize as hospitalization criteria varied from country to country, so these were not included in the learning and test sets. The retrieved sequences contained a total of 3,779 protein and 36 UTR mutation types as compared to the Wuhan strain.

### Association of mutations with patient outcome

We represented the genomes with mutation signature vectors that contained the name of the genome, the age of the patient, followed by a series of protein and UTR mutations listed in the order of their sequence positions.

Our goal was to establish whether or not the mutation signatures can distinguish two input classes, i.e. mild and severe patient outcomes. Machine learning algorithms can help to approach this problem, since a high classification efficiency is generally considered an indicator of the input data being able to distinguish the class labels. A specific problem of the genomic data is the large number of possible mutations which can obscure the identity of truly relevant mutations. Techniques of feature selection (13), (16) are designed to solve this problem as they can narrow down the number of input dimensions to a few, relevant dimensions ranked according to their importance. In practice, we can combine feature selection algorithms, such as LASSO (16) or Statistically Equivalent Signatures (13) with classifier algorithms (such as support vector machines (7,8), random forests (9), logistic regression (10), in such a way that classification performance and feature (mutation) signatures will be optimized at the same time. **Table 1** summarizes the results in terms of classification performance. The AUC and accuracy values (for a review see (12)) are high enough to indicate that the mutation signatures contain the information necessary to separate mild and severe outcomes. **Table 2** lists examples of mutation signatures identified by the LASSO algorithm on two datasets. Here we also included models that did not contain patient age data (which sum up the total of 4 methods listed in **Table 2**). It is conspicuous, that the mutation lists are similar, i.e. almost the same mutations were found to be important in all cases. For instance, out of the 26 different mutations 19 were prioritized by all four methods, four were selected in two of them and there were only three mutations that were found by one method only. Similar if not identical mutation signatures were found with the Statistically Equivalent Signatures method (13) (data not shown). In other words, the prioritized mutations can be considered a stable, robust subset that apparently contain most of the information necessary to distinguish the „Mild” and „Severe” cases. We also note that the mutations prioritized here by feature selection quite well coincide with those well-known from previous sequencing studies. For instance, spike protein variants V1176F and S477N, that co-occur with DG14G, affect important functional domains of the spike protein, and were found to increasingly spread around the world (6). In the nucleocapsid protein, S194L maps onto the phosphorylated “RS-motif” (17) which is in the intrinsically unstructured serine rich region 181-213 of the protein and was previously found associated with severe outcome (5). Similarly, UTR mutations SC241T and SG29830T were also noted by Mukherjee and Goswami (18).

**Table 1.** Prediction classification performance of different methods determined using a balanced dataset of 797 mild and 797 severe genomes (Dataset 1) and 638 mild and 638 severe samples (Dataset 2), using repeated, stratified 10-fold cross-validation, including patient age data. The models used the mutations as well as patient age as the input. The standard deviation of the values was typically <0.02.

|  | Methods<sup>1</sup> |  | Dataset 1 | Dataset 2 |
| --- | --- | --- | --- | --- |
|  | Classification | Feature selection | AUC | AUC |
| 1 | Random Forests | LASSO | **0.943** | 0.936 |
| 2 | Ridge Logistic Regression | LASSO | 0.940 | 0.935 |
| 3 | Random Forest | SES | 0.937 | 0.927 |
| 4 | Ridge Logistic Regression | SES | 0.935 | 0.923 |
| 5 | Support Vector Machine | LASSO | 0.932 | 0.919 |
| 6 | Support Vector Machine | SES | 0.918 | 0.913 |
| 7 | Trivial model | NONE | 0.5 | 0.5 |

<sup>1</sup>The run parameters were as follows: Random Forests: 100 trees with Deviance splitting criterion, minimum leaf size = 3, and variables to split = 1.0 sqrt (nvars); Support Vector machines: type C-SVC with Radial Basis Function Kernel and hyper-parameters: cost = 1.0, gamma = 1.0. LASSO: penalty=1; lambda=2.629e-03; Ridge Logistic Regression: lambda=1; Statistically Equivalent Signatures (SES) (maxK = 2, and alpha = 0.05). The shaded combination is implemented in the online analysis platform. AUC= area under the curve

**Table 2.** Mutation signature examples selected by the LASSO algorithm (16).

|  | 1 | 2 | 3 | 4 |
| --- | --- | --- | --- | --- |
|  | Dataset 1 (incl. age)<sup>1</sup> | Dataset 2 | Dataset 2 (incl. age) | Dataset 2 |
| **Performance<sup>2</sup>:** | AUC=0.938;<br>ACC=0.866 | AUC=0.887;<br>ACC=0.800 | AUC=0.933;<br>ACC=0.836 | AUC=0.893;<br>ACC=0.806 |
| **1** | Spike_V1176F | Spike_V1176F | Spike_V1176F | Spike_V1176F |
| **2** | N_I292T | N_I292T | NS3_Q57H | NS3_Q57H |
| **3** | NS3_Q57H | Spike_L5F | N_I292T | NSP4_M324I |
| **4** | SG29830T | NSP3_A994D | N_D377Y | NSP12_P323L |
| **5** | SC241T | NS3_Q57H | N_S194L | Spike_D614G |
| **6** | N_D377Y | NSP14_A320V | Spike_D614G | NSP13_S485L |
| **7** | NSP4_F308Y | N_S194L | NSP13_H290Y | N_I292T |
| **8** | NS3_G251S | NSP6_L37F | N_G204R | Spike_L5F |
| **9** | NSP6_L37F | SC241T* | NS3_G251V | NSP4_F308Y |
| **10** | NSP4_M324I | N_G204R | NSP14_A320V | N_S194L |
| **11** | N_M234I | SG29830T* | N_M234I | N_G204R |
| **12** | NSP13_H290Y | Spike_D614G | NSP4_F308Y | NSP3_A994D |
| **13** | NSP14_A320V | NSP13_S485L | SG29830T* | NSP6_L37F |
| **14** | N_S194L | NSP4_F308Y | NSP12_P323L | SC241T* |
| **15** | NSP7_S25L | NSP4_M324I | Spike_L5F | NSP3_K945N |
| **16** |  | N_D377Y | NSP6_L37F | NSP7_S25L |
| **17** |  | NSP7_S25L | NSP4_M324I | NS3_G251S |
| **18** |  | NS3_G251V | SC241T* | SG29830T* |
| **19** |  | N_M234I | NS8_L84S | NS3_G251V |
| **20** |  | NS3_G251S |  | NSP14_A320V |
| **21** |  |  |  | NSP13_H290Y |
| **22** |  |  |  | N_D377Y |
| **23** |  |  |  | Spike_N439K |
| **24** |  |  |  | NS8_L84S |
| **25** |  |  |  | NSP3_I1683T |

<sup>1</sup>Determined on datasets described in Methods. The learning procedure only included patient age if indicated.
<sup>2</sup>Corrected AUC values
\*UTR mutations

### Online analysis platform

The complete analysis pipeline is summarized in **Figure 1**. In the first step of the analysis, global pairwise sequence alignment is used to align the query nucleotide sequence to the reference nucleotide sequence (hCoV-19/Wuhan/WIV04/2019) using the “Biostrings” R Bioconductor package (https://bioconductor.org/packages/Biostrings/). A quality control is carried out at this point, and input sequences containing to few identities, too many “N”-s, abnormal GC content, or having a discrepant length with respect to the reference sequence are rejected. Then, using the *translate()* function of the “Biostrings” package, nucleotide alterations are translated to protein alterations plus UTR alterations. The resulting mutation signature is passed on to random forest based predictor that contains a model with patient age. The output is a severity score, which is a (0,1) probability of the infection being severe. This value can be evaluated in comparison with distribution data of the test set **(Figure 2)**. In this figure one can designate approximate segments depending on the ratio of severe and mild outcomes. Namely, score<0.20 and score>0.80 are regions of high confidence, 0.20<score<0.40 and 0.60<score<0.80 are of medium confidence and 0.40<score<0.60 is of low confidence and annotated as “undecided”. An output example is „Score = 0.10, interpretation: mild outcome (high confidence)”, or Score = 0.58, interpretation: “undecided, (low confidence)”. The server contains an option to submit multiple genomes.

**Figure 2.**
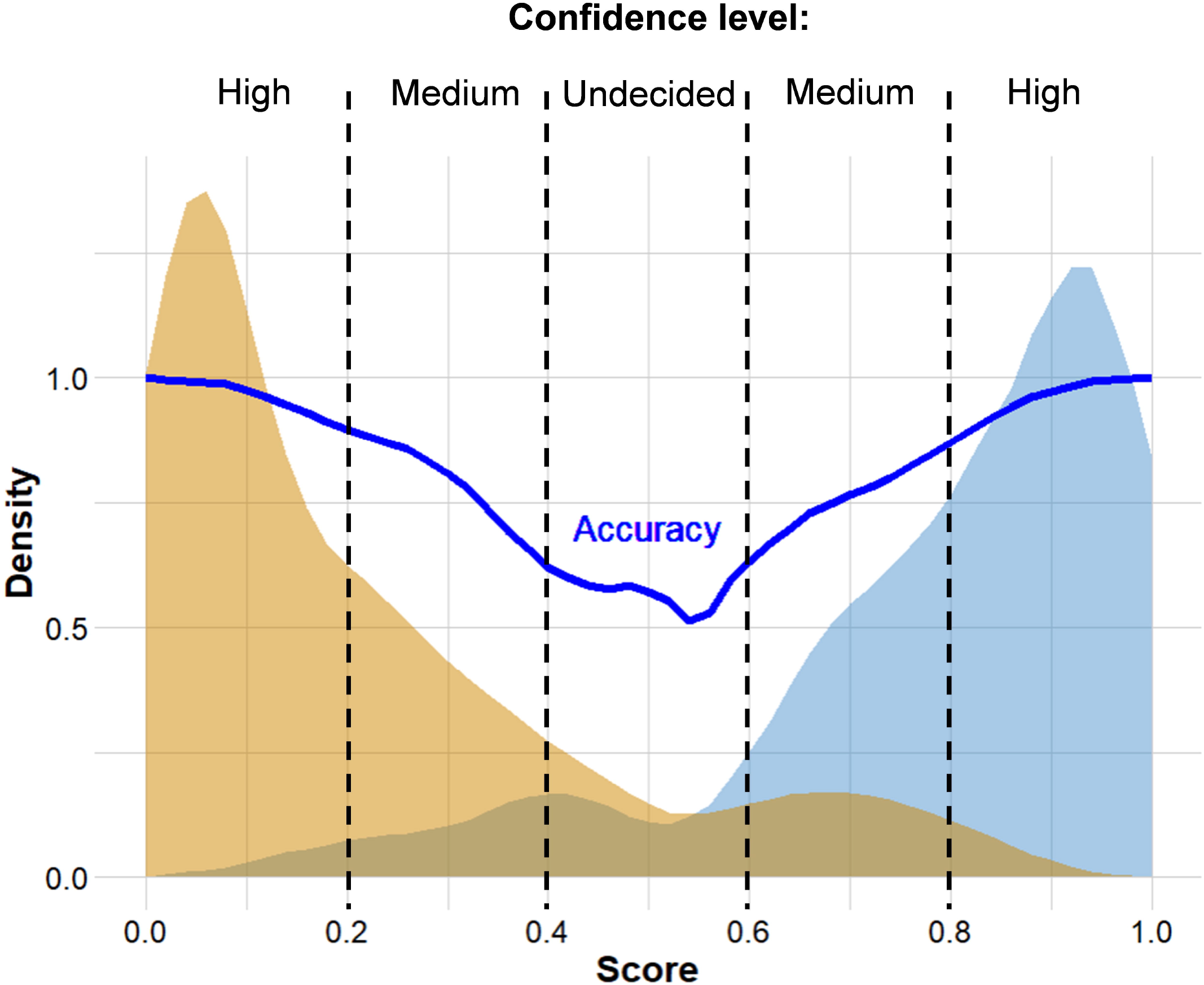
Distribution of scores predicted for genomes associated with known “Mild” and “Severe” clinical outcomes. The thick continuous line indicates confidence defined as the probability of correct prediction, scores below 0.20 and above 0.80 indicate high confidence in predicting “Mild” and “Severe” outcomes, respectively. Intermittent scores are considered medium or low confidence, respectively.

## DISCUSSION

In this work we used machine learning techniques to select mutation signatures associated with severe SARS-CoV-2 infections. We grouped patients into 2 major categories (“mild” and “severe”) by grouping the 179 outcome designations in the GISAID database. A protocol combined of logistic regression and feature selection algorithms revealed that mutation signatures of abouttwenty mutations can be used to separate the two groups. The mutation signature is in good agreement with the variants well known from previous genome sequencing studies, including Spike protein variants V1176F and S477N that co-occur with DG14G mutations and account for a large proportion of fast spreading SARS-CoV-2 variants (6). UTR mutations were also selected as part of the best mutation signatures. The mutations identified here are also part of previous, statistically derived mutation profiles (5)(18).

An online prediction platform was set up that can assign a probabilistic measure of infection severity to SARS-CoV-2 sequences, including a qualitative index of the strength of the diagnosis. The data confirm that machine learning methods can be conveniently used to select genomic mutations associated with disease severity, but one has to be cautious that such statistical associations – like common sequence signatures, or marker fingerprints in general – are by no means causal relations, unless confirmed by experiments.

Our plans are to update the predictions server in regular time intervals. While this project was underway more than 100 thousand sequences were deposited in public databases, and importantly, new variants emerged in the UK and in South Africa that are not yet included in the current datasets. Also, in addition to mutations, we plan to include also insertions and deletions which will hopefully further improve the predictive power of the server.

In summary, we found that automated machine learning, such as the method of Tsamardinos and coworkers used here (15), is a versatile and effective tool to find salient features in large and noisy databases, such as the fast growing collection of SARS-CoV-2 genomes.

## ACKNOWLEDGEMENTS

The research was financed by the 2018-2.1.17-TET-KR-00001 grant and by the Higher Education Institutional Excellence Programme (2020-4.1.1.-TKP2020) of the Ministry for Innovation and Technology (MIT) in Hungary, within the framework of the Bionic thematic programme of the Semmelweis University as well as by OTKA grant 12065 provided by MIT, Hungary to Pázmány University. The authors wish to acknowledge the support of ELIXIR Hungary (www.elixir-hungary.org) and Dr. I. Tsamardinos (University of Crete, Greece) for help and advice.

## SUPPLEMENTAL MATERIALS

**Supplemental Table 1**. Dataset #1 including 797 severe and 797 mild cases.

**Supplemental Table 2**. Dataset #2 including 638 severe and 638 mild cases.

**Supplemental Table 3**. List of database terms mapped to “mild” and “severe” labels in our datasets.

